# Replication stress and FOXM1 drive radiation induced genomic instability and cell transformation

**DOI:** 10.1101/2020.06.29.177352

**Authors:** Zhentian Li, David S. Yu, Paul W. Doetsch, Erica Werner

## Abstract

In contrast to the vast majority of research that has focused on the immediate effects of ionizing radiation, this work concentrates on the molecular mechanism driving delayed effects that emerge in the progeny of the exposed cells. We employed functional protein arrays to identify molecular changes induced in a human bronchial epithelial cell line (HBEC3-KT) and osteosarcoma cell line (U2OS) and evaluated their impact on outcomes associated with radiation induced genomic instability (RIGI) at day 5 and 7 post-exposure to a 2Gy X-ray dose, which revealed replication stress in the context of increased FOXM1 expression. Irradiated cells had reduced DNA replication rate detected by the DNA fiber assay and increased DNA resection detected by RPA foci and phosphorylation. Irradiated cells increased utilization of homologous recombination-dependent repair detected by a gene conversion assay and DNA damage at mitosis reflected by RPA positive chromosomal bridges, micronuclei formation and 53BP1 positive bodies in G1, all known outcomes of replication stress. Interference with the function of FOXM1, a transcription factor widely expressed in cancer, employing an aptamer, decreased radiation-induced micronuclei formation and cell transformation while plasmid-driven overexpression of FOXM1b was sufficient to induce replication stress, micronuclei formation and cell transformation.

## INTRODUCTION

Ionizing radiation is an effective and widely used tool for cancer treatment and control. Over 50% of cancer patients will be exposed to ionizing radiation at some point of their illness (1). Therefore, because of radiation’s wide use and improved cancer treatment outcomes, adverse effects such as malignancies secondary to radiation are becoming more concerning (2, 3). The mechanism for radiogenic cancers is unknown. All tissues are susceptible to develop radiation-induced tumors of low mutational load, without a unique signature resulting from a known mechanism (4, 5). Ionizing radiation is considered a weak mutagen (2, 6), and generates multiple types of lesions on DNA (7). Among them, double strand breaks (DSB) are the most toxic, however can be readily repaired in normal cells or cause death or senescence in repair-impaired cells (8).

Ionizing radiation also generates responses in cells that have not been directly targeted, which include delayed genomic instability, bystander, clastogenic and transgenerational effects (reviewed in (9, 10)). The molecular mechanisms driving these responses and their impact on the overall effects of ionizing radiation remain poorly understood.

Among non-targeted effects, radiation induced genomic instability (RIGI) is a quite heterogeneous response defined by the increased rate of acquisition of *de novo* genomic alterations in the progeny of irradiated cells (11). A diverse set of biological end points have been associated with genomic instability, including micronuclei formation, sister chromatid exchanges, chromosomal gaps, karyotypic abnormalities, microsatellite instability, homologous recombination, gene mutation and amplification, cellular transformation, clonal heterogeneity and delayed reproductive cell death (11). RIGI can be detected following exposure to moderate doses of X-rays in multiple cell lines *in vitro* and in tissues of animals irradiated *in vivo* (9).

Genomic instability is present in the majority of cancers, driving tumor development and evolution. During tumor initiation, genomic instability has been proposed to accelerate the acquisition of mutations. There is evidence supporting the role of genomic instability as a first step in the genesis of certain radiation-induced cancers *in vivo* (6, 12). Thus, RIGI can be considered as a model to study inducible genomic instability and as a mechanism contributing to radiation-induced carcinogenesis (9).

Several factors have been shown to modulate RIGI. Oxidative stress associated with mitochondrial malfunction promotes a genomic instability state, as interference with reactive oxygen species generation or scavenging attenuates readouts (13). Other factors that have been shown to influence RIGI are epigenetic changes and deficiencies in DNA repair (14) (15). However, no unifying molecular mechanism has been established for RIGI development yet.

In the current study we employ an immortalized, non-tumorigenic cell line of human bronchial epithelial cells HBEC3-KT, which have precursor properties as they can differentiate *in vitro* and can be progressed to further transformed states by oncogene expression or exposure to carcinogenic agents (16–18). The lung is an organ remarkably susceptible to low doses of radiation (2, 19) and it is at high risk for carcinogenesis secondary to radiation therapy (20). HBEC3-KT reproduce several of the attributes and end points associated with RIGI. Our previous work showed that exposure to high and low LET radiation induced transient genomic instability peaking at day 7 and decreasing by days 14 and 21 post-irradiation (21) (22). Phenotypes associated with genomic instability such as micronucleus formation and γH2AX and 53BP1 foci are cell autonomous at day 7. Micronucleus formation increases with dose, reaching saturation at 4Gy(23). For the experiments in this work, cells were exposed to a 2Gy dose representing the average radiation dose per fraction received by patients during radiation therapy. Employing protein arrays, we identify two potential mechanisms, replication stress and FOXM1, and evaluate their role in promoting RIGI and cell transformation.

## RESULTS

We have previously shown that radiation induces transient genomic instability in HBEC3-KT cells expressed as increased micronuclei formation, residual 53BP1 and γH2AX foci, in the context of cellular oxidative stress (21). To identify conserved mechanisms driving these phenotypes we leveraged RIGI development in two cell lines of different tissue origin, HBEC3-KT and the osteosarcoma epithelial cell line U2OS to exclude tissue specific mechanism. As shown in Fig. 1A, following exposure to a 2Gy X-ray dose, a three-fold increase in micronucleus formation can be detected at day 7 in HBEC3-KT and at day 5 in U2OS, timing that corresponds to about 4 population doublings. The micronucleus assay detects fragments or whole chromosomes that can not engage with the mitotic spindle as a result of damage to chromosomes or the mitotic machinery (reviewed by M. Fenech (24)). Consistent with damaged DNA at the moment of mitosis, we measured four and two-fold increased frequency of Replication Protein A (RPA) positive chromosomal bridges in the irradiated cell populations for HBEC3-KT and U2OS respectively, an indication of single stranded DNA generation during genomic material segregation (Fig. 1B). An additional reported consequence of DNA damage during mitosis is that 53BP1 bodies accumulate in the following G1 phase (25). Both cell types showed roughly a three-fold increased accumulation of 53BP1 bodies in cells in G1 phase, identified by the absence of nuclear Cyclin A (Fig. 1C). RPA positive chromosomal bridges have been previously observed in cells that undergo mitosis in the presence of accumulated unresolved homologous recombination intermediaries (26, 27). To test whether the homologous recombination repair pathway is used at a higher frequency in irradiated cells, we utilized a panel of genetically engineered U2OS cell lines with integrated GFP constructs to report the repair of a chromosomal DNA double strand break introduced by the endonuclease I-SceI by homologous recombination (HR), non-homologous end joining (c-NHEJ) and alternative non-homologous end joining (a-NHEJ) (Fig. 1D, Supplementary Fig.1)(28). Compared to non-irradiated cells, exposure to X-ray increased the number of GFP positive cells detected in the HR reporter line by 2.5 fold while the relative number of cells detected decreased in the c-NHEJ reporter line without affecting a-NHEJ dependent repair. Increased HR is a previously described RIGI outcome (29). Altogether, these results suggest that several days following irradiation, replicating cells are undergoing mitosis with damaged DNA and relying more on the HR pathway to repair double strand breaks.

**Figure 1:**
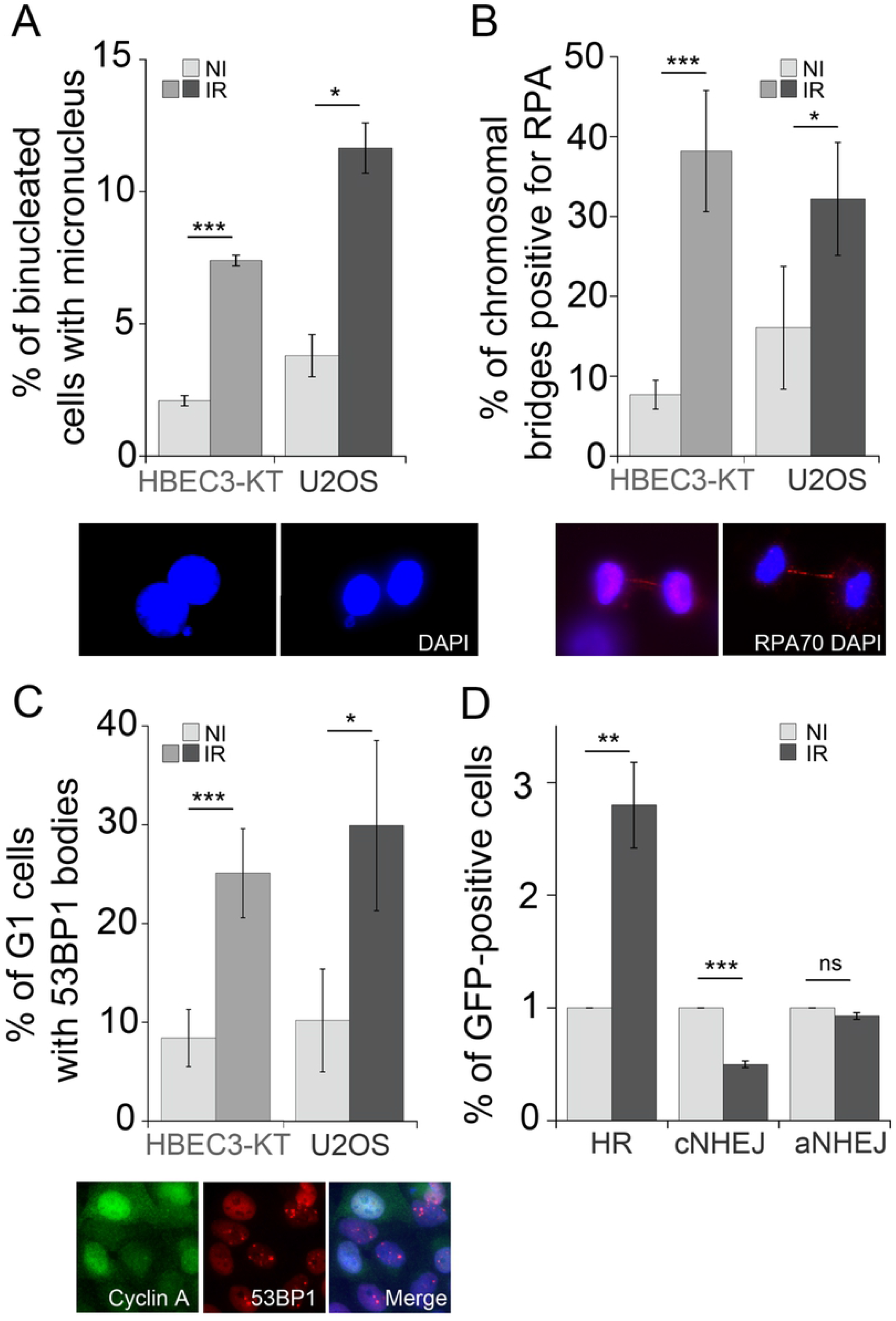
X-ray irradiated cells have damaged DNA at mitosis and increased usage of the HR pathway. A) Micronucleus formation rates at day 7 (HBEC3-KT) or day 5 (U2OS) following exposure to 2Gy X-ray. Average of 2 experiments, error bars represent SEM. Student’s t-test. Inserts depict a representative image of binucleated HBEC3-KT cells with a micronucleus or a nuclear bud, respectively. B) Increased frequency of RPA-positive chromosomal bridges in irradiated cells. Proliferating cultures of day 7 (HBEC3-KT) or day 5 (U2OS) following exposure to 2Gy X-ray were fixed and stained for RPA 70. An average of 90 mitosis per condition were scored for RPA positive bridges. Error bars represent standard deviation of triplicate samples. Student’s t-test. Insert depicts a representative image of RPA-positive chromosomal bridge in U2OS (left panel) and HBEC3-KT (right panel) cells. C) Frequency of nuclei with more than three 53BP1-positive foci per nuclei scored in an average of 100 Cyclin A negative nuclei per sample in HBEC3-KT at day 7 and in U2OS cells at day 5 following a 2Gy X-ray dose. Error bars represent standard deviation of triplicate samples. Student’s t-test. Insert depicts HBEC3-KT cells with Cyclin A positive nuclei without 53BP1 bodies and Cyclin A negative nuclei with 53BP1 bodies. D) Relative GFP induction levels in U2OS reporter cell lines for homologous recombination dependent repair (DRG), cNHEJ (EJ2) and aNHEJ (EJ5) at day 7 following exposure to a 2Gy X-ray dose. Average of 2 experiments, error bars represent SEM.

To gain further understanding of the mechanisms and scope of the alterations occurring in irradiated cells, we performed a functional proteomics analysis employing reverse phase protein arrays (RPPA) to assess the expression of 431 proteins including post-translational modifications relevant to cancer. Furthermore, this array is enriched in proteins involved in cell cycle regulation and HR. We analyzed the responses to a 2Gy dose X-ray at day 5 and 7 of HBEC3-KT cells and at day 5 for U2OS. Unsupervised hierarchical clustering of the results for HBEC3-KT show a clear segregation of the day 7 samples from the non irradiated and day 5 post-irradiation samples (Supplementary Fig. 2), which is consistent with our previous findings that the phenotype at day 7 is a response that emerges overtime after exposure and not residual response to initial damage(22). We reasoned that proteins and pathways that co-variate in both cell lines could be associated with common phenotypes induced by radiation exposure. A comparison between the proteins significantly differentially regulated in U2OS and HBEC3-KT (Fig. 2A) reveals common up-regulation of phospho Aurora Kinase A and an increase in FOXM1 expression, a transcription factor that regulates the expression of multiple DNA repair proteins, increases HR dependent repair in the direct repeats conversion assay (30, 31) and cell cycle progression (32). Analysis focused on proteins involved in cell cycle regulation and HR is shown in Fig. 2B. The heat map exhibits an overall milder response for U2OS than HBEC3-KT cells, but both cell models display increased expression of markers related to end-resection (CtIP in U2OS and S4/S8 RPA phosphorylation in HBEC3-KT) in the context of small changes in levels and phosphorylation of proteins involved in checkpoint activation such as ATM, ATR, Chk1 and Chk2. The interrogation also reveals upregulation of proteins promoting M phase entry reflected by increases in CDK1 and Wee1 phosphorylation with increased PLK1 and Cdc25C, which are more prevalent in HBEC3-KT, but supported by a significant decrease in p21 expression in U2OS cells. In HBEC3-KT, RAD51 and Cyclin B, two targets for FOXM1 regulation, had increased expression at day 7. Collectively, these findings suggest the presence of a modest replication stress and checkpoint activation within the context of increased FOXM1 expression and activity.

**Figure 2:**
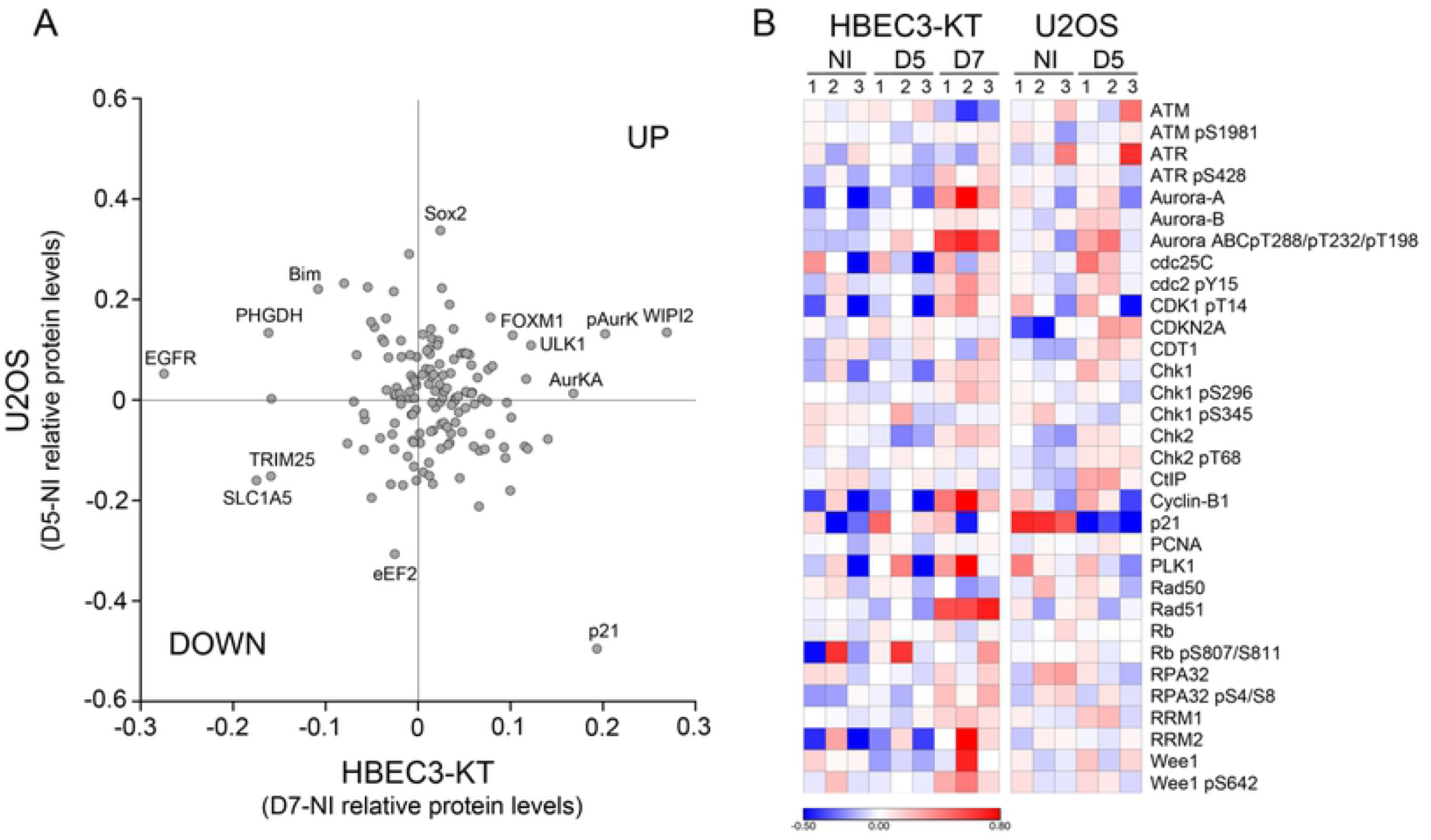
Reverse Phase Protein Array analysis of proteins and post-translational modifications altered by exposure to X-ray. A) Scatterplot representing the relative protein level difference between irradiated and non-irradiated samples. Averages of triplicate samples normalized and transformed to linear values for each condition were subtracted. Each dot represents a protein that yielded a significant difference after irradiation (Student’s t-test, p<0.05) in cell lysates of HBEC3-KT at day 7 and/or U2OS at day 5. B) Heat map for proteins involved in cell cycle regulation and homologous recombination DNA repair pathway. The normalized log2 values for protein levels in each sample were median centered for each protein measured.

To validate these findings, we measured DNA replication rate employing the DNA fiber assay. As shown in Fig. 3A, replication speed was significantly slower (all doses vs. non irradiated p<0.0001, one way ANOVA) in cells that have been irradiated, reduced from 0.8 kb/min in non-irradiated cells to 0.61, 0.54 and 0.59 following doses of 2, 4 or 6 Gy. As the radiation dose increases, a larger fraction of the replication tracks become shorter, but the assay did not detect pausing or stalling as all tracks incorporated both labeled nucleotides and no significant track labeling asymmetry was detected (Supplementary Fig. 3A) (33). As a reference, we measured a replication rate of 0.26 kb/min in cells treated with 25μM hydroxyurea for 24h. This low dose has been reported to cause replication stress by reducing the availability of nucleotides following inhibition of ribonucleotide reductase (34). Reduced replication speed leads to increased single stranded DNA accumulation, which binds RPA and commits damage repair by HR. This prediction was examined by immunofluorescence to detect RPA associated with chromatin, which revealed that exposure to radiation increased by three-fold the percentage of cells with RPA foci in HBEC3-KT at day 7 and in U2OS at day 5 (Fig. 3B). These results are consistent with increased RPA phosphorylation detected by RPPA (Fig. 2B). Reduced availability of nucleotides is a mechanism leading to replication stress frequently found in cancer and can be reversed by nucleoside supplementation (35). However, addition of nucleosides to the cultures in conditions that reduced micronucleus formation induced by HU was not effective in reducing radiation-induced effects (Supplementary Fig. 3B). FOXM1 is one of the proteins significantly elevated at day 7 but not at day 5 post-irradiation in HBEC3-KT cells and increased in U2OS cells as well (Fig. 3C), which we confirmed by western blot in lysates from both cell lines (Fig. 3D). Interestingly, FOXM1 expression as well as RAD51 and EXO1, two genes that have been reported to be transcriptional targets for FOXM1 (36, 37) also increased in HBEC3-KT at day 7. U2OS cells have a higher basal FOXM1 expression, which was increased by radiation, but the up-regulation of the transcriptional targets measured was less robust than in HBEC3-KT cells.

**Figure 3:**
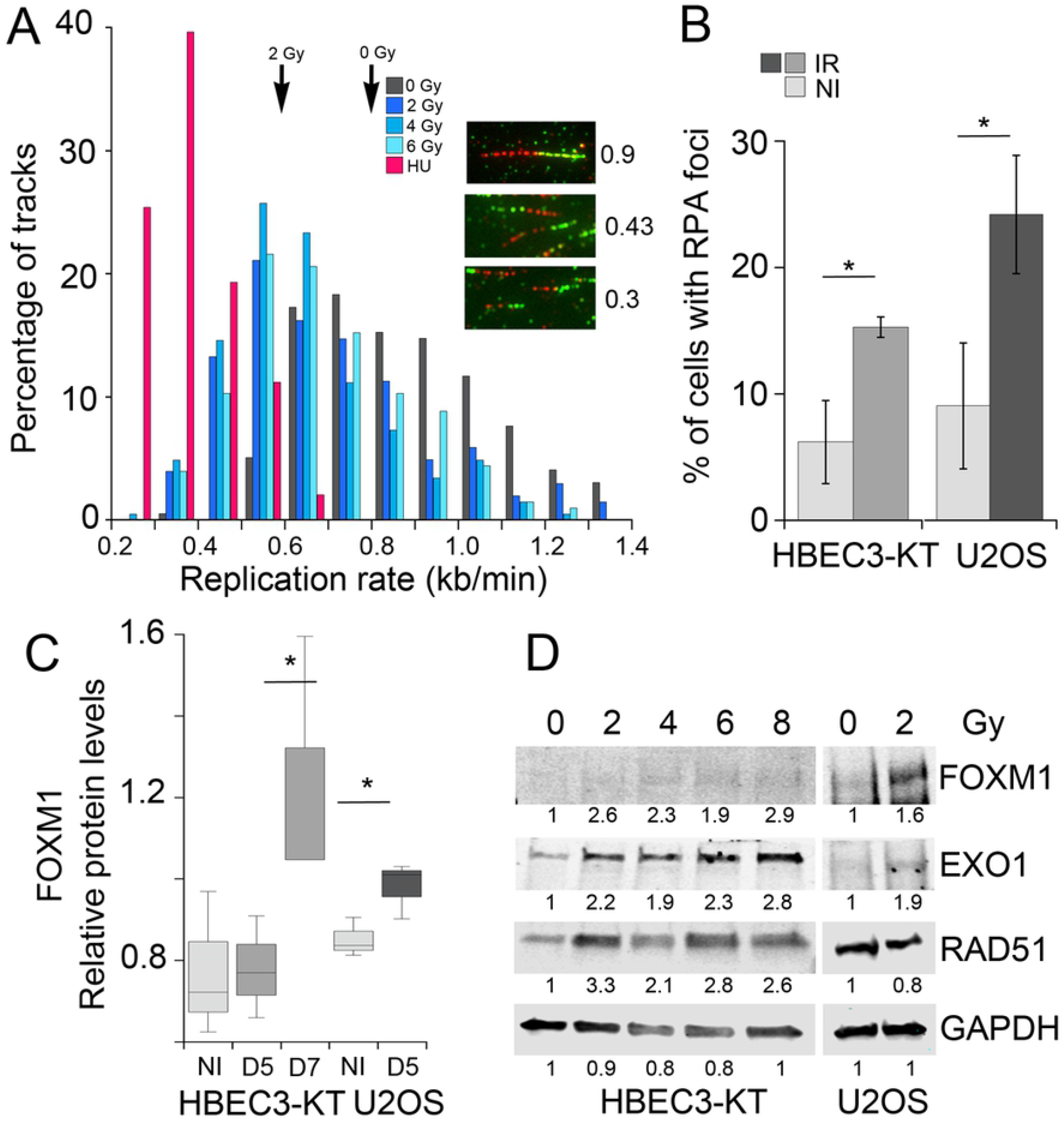
Irradiated cells have low levels of replication stress and induce FOXM1 expression. A) Frequency distribution plot of replication speed measured by the DNA fiber assay in HBEC3-KT cells at day 7 following exposure to the indicated X-ray doses. As a control, HBEC3-KT cells were treated for 48h with 25μM HU to reduce replication speed. Arrows indicate average speed. Inserts show examples of replication tracks of different length. B) Quantification of the percentage of cells with RPA70 foci detected by immunofluorescence. Between 50 and 100 cells were scored in replicate samples of non-irradiated and day 7 irradiated HBEC3-KT cells or day 5 irradiated U2OS cells. Error bars represent standard deviation. Student’s t-test. C) Boxplots for the relative FOXM1 expression levels detected by reverse phase protein arrays in HBEC3-KT at day 5 and 7, and in U2OS at day 5 post-exposure to a 2Gy X-ray dose. Student’s t-test. D) Western blot for FOXM1 expression and known transcriptional target proteins in HBEK3-KT at day 7 and in U2OS at day 5 post-exposure to the indicated X-ray dose.

Given the significant impact that FOXM1 expression has on the expression of DNA repair proteins and on carcinogenesis, we tested whether it is contributing to the genomic instability phenotypes in irradiated cells employing an aptamer oligonucleotide that was selected to interfere with transcriptional activity of FOXM1 (Aptamer) and compared to an oligomer of random sequence (Control) (38) instead of FOXM1 knockdown, because it has been shown that reduction in FOXM1 protein disrupts mitosis and causes DNA damage (39, 40).The FOXM1-specific aptamer reduced the radiation-induced upregulation of EXO1, Cyclin B and RAD51 when delivered by transfection to HBEC3-KT cells (Fig. 4A) without increasing micronucleus (Figs. 4B, 6B) or HR (Fig. 4C) in non-irradiated cells. The aptamer significantly reduced micronucleus formation rates in irradiated HBEC3-KT cells by 50% (Fig. 4B), and reduced the number of GFP-positive cells resulting from homology-dependent repair in both irradiated and non-irradiated U2OS cells (Fig. 4C). Taken together, these results suggest a role for FOXM1 in promoting RIGI in both cell types.

**Figure 4:**
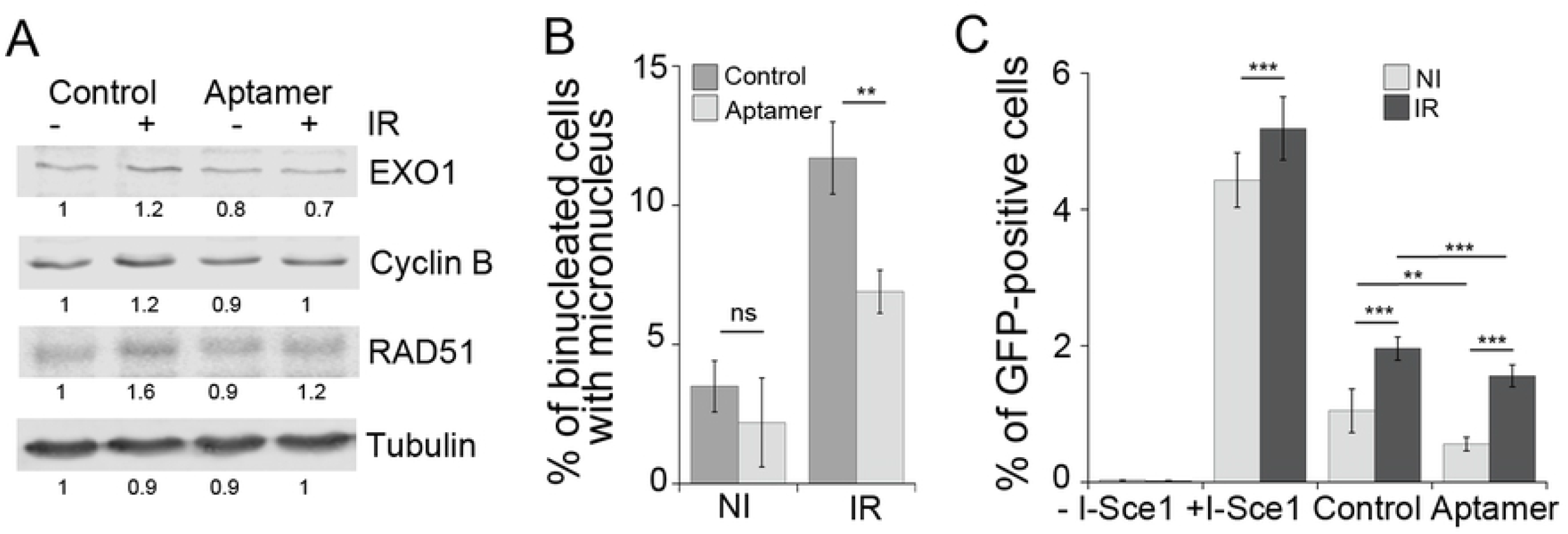
Interference with FOXM1 transcriptional activity reduces radiation induced phenotypes. A) Western blot analysis of known transcriptional target proteins for FOXM1 in HBEC3-KT at day 7 following exposure to 2Gy X-ray and transfection at day 3 with 100nM aptamer of random sequence (Control) or of a sequence interfering with FOXM1 transcriptional activity (Aptamer). B) Micronucleus assay in the same cells analyzed in A. Error bars represent standard deviation. 1 of 2 experiments shown. One Way ANOVA followed by Bonferroni post-test. C) Effect of aptamer transfection on the gene conversion assay to report homologous recombination dependent repair. Error bars represent SEM. Average of two experiments. One Way ANOVA followed by Bonferroni post-test.

**Figure 5:**
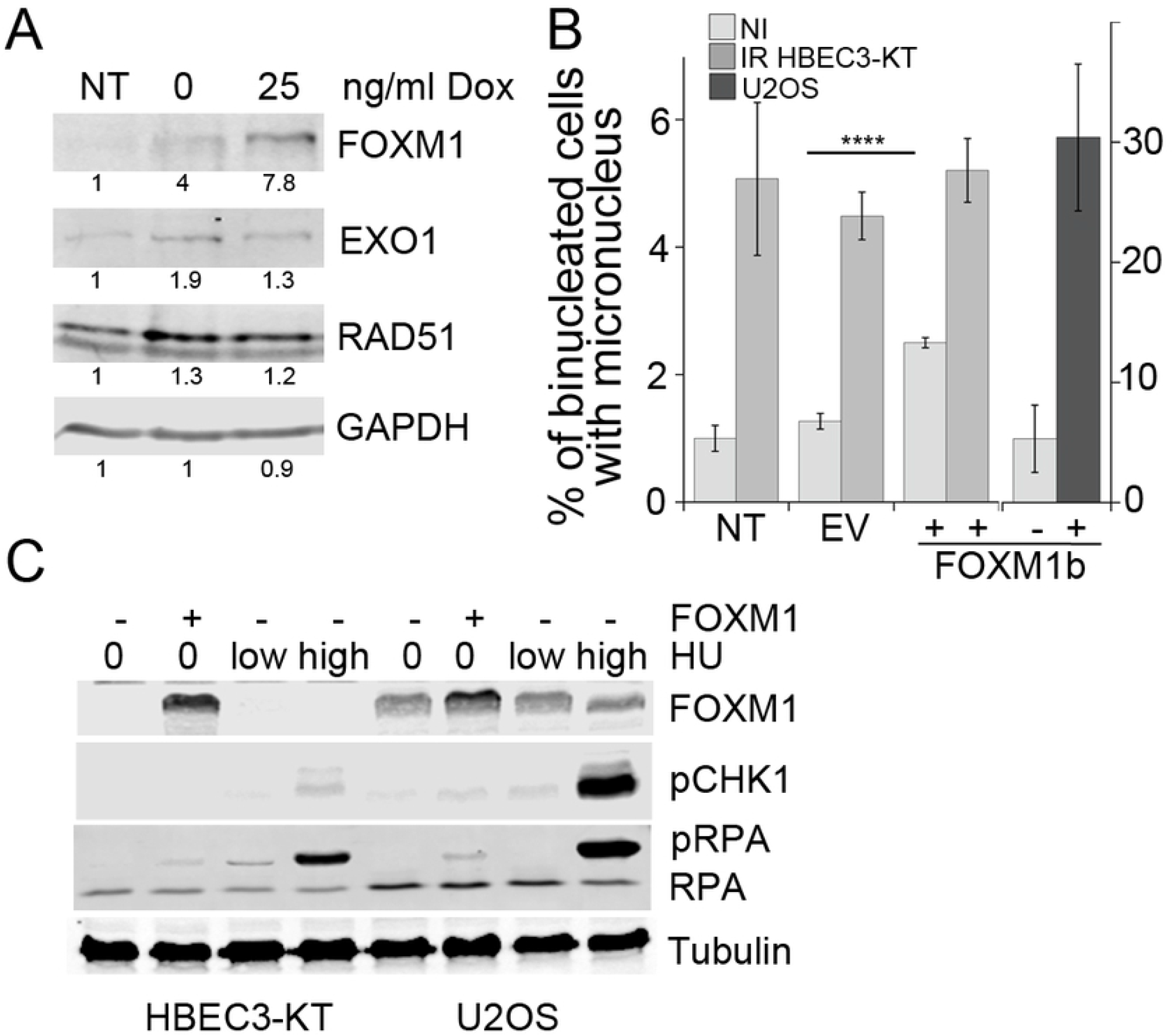
FOXM1b overexpression is sufficient to induce genomic instability. A) Western blot for Flag-tagged FOXM1 expression and transcriptional target proteins in stably expressing U2OS cells induced for 48h with the indicated Doxocyclin concentration. B) Micronucleus formation in HBEC3-KT and U2OS transiently expressing FOXM1. Empty vector or FOXM1 were transfected 72h prior to the 18h incubation with cytochalasin. U2OS cells were treated with 1μg/ml Dox. Error bars represent SEM. Student’ t-test. C) Western blot analysis for RPA2 and Chk1 phosphorylation in HBEC3-KT and U2OS cells overexpressing FOXM1 or treated with HU 25μM for 48h or 3mM for 4h.

**Figure 6:**
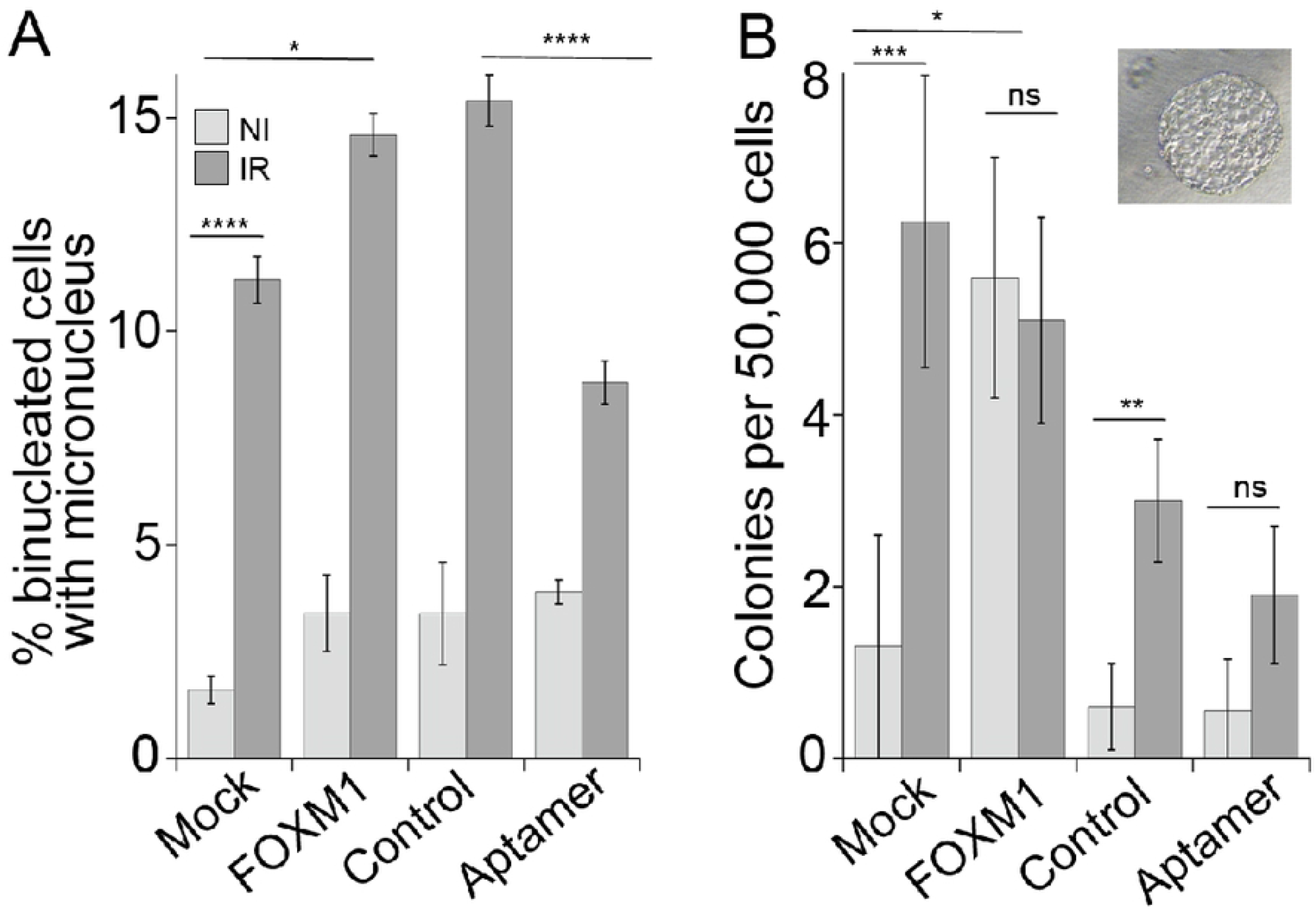
FOXM1 expression promotes cell transformation. Triplicate flasks of HBEC3-KT cells were exposed to 4Gy X-ray and transfected at day 4 with the indicated construct. At day 6 post-irradiation, the cells were passaged and samples were collected for micronucleus assay shown in (A). After a month of continuous growth, the cells were tested for growth in soft agar shown in (B). Error bars represent SEM. Student’ t-test and One Way ANOVA with Fisher’s LSD post-test for aptamer and radiation samples.

We tested next, whether FOXM1 overexpression is sufficient to induce some of the radiation-induced phenotypes. To increase FOXM1 expression levels we employed a cDNA construct regulated by Doxocyclin (Fig. 5A), which in U2OS cells increased the expression of EXO1 and RAD51 under our cell culture conditions even without Doxocyclin addition. FOXM1 overexpression, but not empty vector, was sufficient to induce micronucleus formation in HBEC3-KT and U2OS (Fig. 5B). FOXM1 overexpression, was sufficient to induce RPA phosphorylation in Serines 4 and 8, similar to the induction produced by a 48h treatment with 25μM HU or a 4h treatment with 3mM HU, conditions that induce mild and high replication stress, respectively (Fig. 5C), consistent with previous reports (34). FOXM1 overexpression also induced mild increases in CHK1phosphorylation, which is an indicator of checkpoint activation occurring as a response to disruptions in S phase. Collectively, these results indicate that FOXM1 expression alone induces replication stress as indicated by RPA and CHK1 phosphorylation and defects during mitosis measured by the micronucleus assay, reproducing some of the phenotypes induced by radiation.

The RPPA platform has been widely employed to analyze cell lines and tumor samples in the TCGA database and the results have been compiled in searchable databases (41, 42). We asked next, how general is the correlation between FOXM1 expression and markers for replication stress (pS345 Chk1 and/or pS428 ATR) and FOXM1 transcriptional targets Cyclin B and RAD51 in cancer cell lines and tumor samples. A survey of the databases revealed that the cell lines derived from multiple cancer types show a significant (p<0.05) positive correlation of FOXM1 with Cyclin B and/or RAD51 expression suggesting FOXM1 activity, which did not necessarily correlate with average FOXM1 expression levels of each cancer type category (Fig. S4A, B). The cell lines of 3 cancer types had a strong (R>0.66) and another 3 had a moderate (0.65>R>0.35) positive correlation between FOXM1 expression and a marker for replication stress. A significant positive correlation, although moderate (0.65>R>0.35) between FOXM1 and pS345 Chk1, is observed in tumor samples from liver, ovarian, cervical and testicular cancer (Supplementary Fig. 5B). In tumor samples, liver, cervical and testicular tumors had higher levels of FOXM1 (Supplementary Fig. 5A). These observations suggest that this is a mechanism highly represented in cancer lines and in a wide range of tumor types.

Given that our results show that radiation and FOXM1 induce replication stress, a known driver of cancer initiation and progression, we tested next whether exposure to radiation promotes cell transformation in HBEC3-KT and involves FOXM1 in this process. HBEC3-KT cells are immortalized but not transformed (16). However, transformation and progression towards malignancy can be induced by exposure to carcinogens such as radiation, and cigarette smoke condensate (43, 44) and by expression of several oncogenes (45). As an intermediate step in transformation and progression, HBEC3-KT cells acquire the capability to form colonies in soft agar, which measures anchorage independent growth. Cell cultures were irradiated in triplicate with a 4 Gy dose to increase the sensitivity of the assay by inducing a greater response measured by reduced DNA replication (Fig. 3A) and higher levels of micronuclei formation (Fig 6A) and (23). At day 7, the effects of the different treatments were evaluated by micronucleus formation shown in Fig. 6A. At 4 weeks of continuous growth, the cells were evaluated for the capability to grow in soft agar. Exposure to radiation significantly increased the number of colonies in mock and in control aptamer transfected cells. Transient interference with FOXM1 activity at the time of genomic instability by the aptamer, reduced the radiation response, demonstrating a role for both, RIGI and FOXM1 activity on cell transformation events. All together these results suggest that low levels of FOXM1 activity, such as the ones induced by radiation exposure, are sufficient to promote cell transformation.

## DISCUSSION

In this work we have uncovered two mechanisms, replication stress and FOXM1 expression, driving RIGI following exposure to X-rays. We employed functional proteomics to evaluate the expression of a selected number of proteins in two cell lines, HBEC3-KT and U2OS, exhibiting RIGI at days 7 and 5 post-irradiation respectively. The signature of changes detected indicated the presence of replication stress and implicated FOXM1 as a candidate molecule to promote RIGI. These predictions were confirmed by measuring slower DNA replication rates with the DNA fiber assay and accumulation of ssDNA in irradiated cells as well as by modifying RIGI outcomes by altering FOXM1 activity. Furthermore, employing these tools to interfere with delayed genomic instability, we demonstrate a contribution of RIGI to radiation-induced cell transformation of HBRC3-KT cells *in vitro*.

One of our most significant finding is replication stress in cells undergoing RIGI, which mechanistically may explain several of the outcomes associated with RIGI measured in previous studies and add novel outcomes to further describe this non-targeted effect. Employing the DNA fiber assay, we detected lower replication elongation rates in irradiated cells, slowing down as the dose of the initial exposure increases (Fig. 3A). Reduced processivity of the replication machinery is known to be sufficient to cause uncoupling with replicative helicases, exposing RPA-binding ssDNA(46), which we detected as increased RPA-positive foci (Fig. 3B). HR (Fig. 1D) is activated as a tolerance mechanism to repair DNA DSBs that are more prone to occur and to stabilize and repair replication forks (47) and is a RIGI outcome following low and high LET radiation exposures (29). An additional consequence of reduced replication speed is the incomplete replication of the genome, particularly in late replicating regions rich in chromosomal fragile sites or of low density in replication origins (48). Non-replicated regions result in nondisjunction of sister chromatids during mitosis, which leads to the formation of anaphase bridges (49), often exposing ssDNA, which we detected as RPA-positive bridges (Fig. 1B). RPA positive chromosomal bridges, associated with chromosomal aberrations, are observed when replication stress is induced in cancer cells (35). DNA damaged during mitosis, is protected by 53BP1 binding until repair during the following phase of the cell cycle (25), which we measured in irradiated cultures as an increased frequency of 53BP1 bodies in Cyclin A-negative cells (Fig. 1C). A screen to identify factors contributing to the formation of 53BP1 bodies found that the most effective condition was a low aphidicolin dose, sufficient to impair replication fork progression through physiological barriers (48), supporting the notion that a reduction in replication elongation rate such as the observed in our work, would be sufficient to induce all these RIGI phenotypes (25). Replication stress could also explain additional RIGI outcomes previously described in the literature but not investigated in this work: Slow or arrested replication forks stimulate the use of potentially mutagenic DNA repair pathways that can lead to the accumulation of mutations and to chromosomal rearrangements (reviewed by Gaillard et. al (46)). Replication stress induces DSB formation and ssDNA gaps which lead to chromosomal fragility and gap formation at metaphase (35). Increased HR activity is associated with increased sister-chromatid exchanges and recombination events (reviewed in references (50–52)).

The main causes for replication stress have been attributed to the reduced availability of resources required for replication, including nucleotides or histones as well as structural impediments in the DNA, such as damage, bound proteins or DNA secondary structures (53). We examined whether a deficiency in the nucleotide pool was a factor, which is a common mechanism leading to replication stress in oncogene-driven cancer (35), however increasing the availability of nucleotides had no effect (Supplementary Fig. 3B). Another factor affecting replication rate could be chronic oxidative stress, which we have previously shown to be present in HBEC3-KT at day 7-post irradiation (21). Chronic oxidative stress has been shown to slow replication in HR defective cells, however this deficiency can be suppressed by supplying nucleotide precursors (54). Low levels of ROS have been shown to induce the dissociation of the component of the replication protection complex Timeless, from the replisome causing fork slowdown (55) and the modification of several other factors involved in replication (53). However, our previous characterization of the oxidative stress response in these cells indicates that oxidative stress is a permissive factor for genomic instability engaged at low dose, while many of the genomic instability readouts show dose dependency pointing to an additional determinant factor (21, 23).

We identified FOXM1 as a second factor modulating RIGI outcomes. FOXM1 is a transcription factor widely expressed in pre-malignant lesions and cancer, and is included in a chromosomal instability signature correlating with functional aneuploidy and highly predictive for clinical outcomes (56, 57). Our findings are consistent with FOXM1 activity promoting genomic instability. We show that radiation increased FOXM1 expression with dose at day 7 post-exposure (Fig. 3C,D) and that FOXM1 overexpression is sufficient to induce RPA and Chk1 phosphorylation (Fig. 5C), considered as the most specific indicators for replication stress (58, 59). While the induced phosphorylation levels were small, they were comparable to low doses of HU, which we show reduced replication rate and increased micronucleus formation (Supplementary Fig. 3B). Low doses of aphidicolin or HU are sufficient to reduce replication speed without significant ATR and Chk1 activation (34, 60) have been also shown to induce 53BP1bodies optimally (25) and expression of CFS (48). Replication stress would offer a mechanism to explain reported FOXM1 dependent increases in LOH, and CNV (61, 62), which have been also identified as RIGI outcomes. Interference with FOXM1 transcriptional activity reduced radiation-induced micronucleus formation and increased HR dependent repair, while overexpression was sufficient to induce micronuclei formation (Figs. 4 and 5).

FOXM1 controls the expression of multiple genes involved in DNA homeostasis including NBS, BRIP1, XRCC1, BRCA2, EXO1, RAD51 and RRM2 as well as cell cycle regulation, Cyclin B and PLK1 ((30, 31, 36, 37, 40, 63, 64) and reviewed in (32)). Several of these targets could mediate FOXM1 role during RIGI. Overexpression of RAD51 increases HR activity and chromosomal instability (65–67), and overexpressed EXO1 promotes end resection (68). An alternative mechanism could be by promotion of premature entry into mitosis by inducing the expression of Cyclin B and PLK1 (39). Given these pleiotropic effects, further experiments beyond the scope of this work, are needed to identify the transcriptional target of FOXM1 promoting RIGI.

We show evidence supporting a role for FOXM1 in radiation-induced cell transformation as transient interference with FOXM1 transcriptional activity was sufficient to dampen cell growth in soft agar induced by IR (Fig. 6B). This activity is consistent with transgenic mice models showing that FOXM1 expression promotes early steps in tumorigenesis, promoting Clara cell and hepatocyte hyperplasia for example (69, 70) and acting in conjunction with many other carcinogens (71) as well as cancer cell lines (72, 73). Replication stress has been shown to be a source for genomic instability in pre-neoplasic lesions, while in normal cells replication stress leads to cell death or senescence through activation of DDR (74–76). FOXM1 expression could be a tolerance mechanism to evade anti-proliferative signals, such as oncogene-induced differentiation (77), DNA damage-induced senescence (31) and aging (78), and allow the proliferation of genomically unstable cells. Thus, we propose that FOXM1 expression is enabling radiation induced cell transformation.

In summary, we show that HBEC3-KT is a model system to elucidate mechanisms driving RIGI, as it reproduces many of the outcomes for RIGI reported in other model systems. Moreover, by transient interference with FOXM1 activity, we demonstrate that the observed transient increase in genomic instability contributes to radiation-induced cell transformation and is thus a potential target of intervention to mitigate late emerging deleterious effects of radiation exposure.

## ACKNOWLEDGMENTS

We thank Kathryn Aziz for coordinating sample analysis at the MD Anderson RPPA core, funded by NCI#CA16672. Flow cytometry was performed in the Emory +Pediatric’s/Winship Flow Cytometry Core.

## AUTHOR CONTRIBUTIONS

Conceptualization E.W.: Investigation, Z.L., E.W.; Resources: D.S.Y.; Writing – Original Draft, Z.L., E.W.; Writing – Review and Editing, Z.L., E.W., D.S.Y, P.W.D.; Funding Acquisition, E.W. and P.W.D.

## ABREVIATIONS

IR: ionizing radiation
LET: Linear Energy Transfer
LOH: loss of heterozygocity
CNV: copy number variation
ssDNA: single stranded DNA
dsDNA: double stranded DNA
CFS: chromosomal fragile site

## MATERIALS AND METHODS

### Cell lines, reagents and irradiations

The human bronchial epithelial cell line (HBEC3-KT) was a gift from Dr. Story (UT Southwestern), was authenticated by karyotyping and tested for mycoplasma at the moment of stock preparation. Cells were cultured in Keratinocyte serum free media (Invitrogen) supplemented with antibiotics, Epidermal growth Factor and Bovine Pituitary extract. HBEC3-KT cells were transfected with Fugene HD or Lipofectamine 3000. FOXM1 and control aptamers were custom synthesized by Sigma with thio-modification, prepared and used at a 100nM concentration as described in (38). DRG, EJ5 and EJ2-U2OS cells were a gift from Dr. Jeremy Stark (Beckman Research Institute of the City of Hope). pCW57.1-FOXM1b was a gift from Adam Karpf (Addgene plasmid # 68811), pRRL sEF1a HA.NLS.Sce (opt).T2A.IFP was a gift from Andrew Scharenberg (Addgene plasmid # 31484).

Low-LET irradiation was carried out using an X-ray machine (X-RAD320, Precision X-Ray, N. Branford, CT, USA) at 320kV, 10mA. Irradiation was delivered at room temperature as a single dose or multiple fractions of 320 kV X-rays (Precision X-Ray Inc., North Branford, CT, USA), at dose-rate approximately 2.3 Gy/min. For irradiations, 200 000 cells were plated in a T25 flask two days before irradiation in triplicates to ensure continuous proliferation before and after exposure. At day 4, each flask was subcultured at a1:3 ratio.

### DNA Repair reporter assay

DRG, EJ5 and EJ2 U2OS reporter cell lines, a gift from Dr. Jeremy Stark, were grown in DMEM 10% FBS. At day 3 after irradiation, 500 000 cells were transfected with I-SceI IFP in triplicate. At day 6, all cells were incubated with 25μM biliverdin, and 18h later the cells were collected to measure expression of GFP and IFP (for transfection efficiency normalization) by flow cytometry in a FACS LSRII Instrument (BD Biosciences). The data collected was analyzed using FlowJo software as described (79). Control and FOXM1 aptamer were transfected at 100nM at day 2 after irradiation.

### Micronucleus assay

Cells were plated at a density of 20 000/well on glass coverslips and treated for 18h with 3 μg/ml Cytochalasin B in media to block cytokinesis before fixing with 4% PFA and staining with DAPI. Binucleated cells, which are the cells that underwent mitosis during the cytochalasin incubation period, were scored for the presence of micronuclei and/or nuclear blebs as previously described (21).

### Reverse Phase Protein Array

Cell lysates were prepared from three biological replicates in 50mM Hepes pH7.4, 1% Triton X-100, 150mMNaCl, 1.5mM MgCl_2_, 1mMEGTA, 10% Glycerol containing anti proteases and anti phosphatases (Roche) and sent for analysis at the Reverse Phase Protein Array Core Facility at MD Anderson. Relative protein levels are obtained by interpolating several sample dilutions into a standard curve and then normalized for protein loading. The scatter plot was generated by graphing the difference between irradiated and non-irradiated averages of triplicate samples normalized and transformed to linear values. Included are only the proteins that yielded a significant difference (paired t-test, p<0.05) in HBEC3-KT and/or U2OS cell lysates. Heat maps were generated with normalized log2 median centered data for each marker employing Morpheus software (Broad Institute).

### DNA Fiber assay

Cells were pulsed with 25μM IdU and then with 250μM CldU for 30 minutes each at 37°C then released with trypsin and spotted on glass slides. DNA was spread following a 7min lysis with 0.5% SDS, 200 mM Tris-HCl pH 7.4, 50 mM EDTA. Dried slides were fixed with a (3:1) Methanol/Acetic acid mixture and denatured for 60min in 2.5 N HCl. Following 1h blocking in 15%FBS in PBS, the slides were incubated in primary antibodies for 2h 1:400 rat anti-bromodeoxyuridine (ab6326 Abcam) and 1:50 mouse anti-bromodeoxyuridine (B44, BD). After a stringency wash with 10 mM Tris–HCl pH 7.4, 400 mM NaCl, 0.2% Nonidet P40 (NP40), the slides were incubated with Alexafluor conjugated secondary antibodies: 488-conjugated chicken anti-rat, 488-conjugated goat anti-chicken, 594-conjugated rabbit anti-mouse and 594-conjugated goat anti-rabbit in 1:300 dilution. The slides were mounted in Fluoromount-G (Southern Biotech). Images of the fibers were acquired with a 63x oil immersion objective on a Zeiss Observer Z1 microscope equipped with Axiovision 4.8 software. 200 dual color labeled replication track length was measured employing Fiji Software and were converted to replication speed using conversion factor is 2.59kb/μm (80, 81). Statistical significance between groups was assessed using One Way ANOVA with a Bonferroni all pairs comparison post-test.

### Immunofluorescence

RPA foci and anaphase bridges were detected in cells fixed with 4% paraformaldehyde and permeabilized with 0.2% TritonX-100. For RPA foci, cells were pre-extracted with 0.5% TritonX-100 before fixing. Antibodies used were RPA70 (Millipore), 53BP1 (Novus Biologicals), Cyclin A (Santa Cruz). The slides were mounted in Fluoromount-G (Southern Biotech). Images were acquired with a 63x oil immersion objective on a Zeiss Observer Z1 microscope equipped with Axiovision 4.8 software. Images were processed using contrast/brightness enhancement only. Foci were counted in at least 5 different fields totaling 50 or more cells in duplicate irradiations.

### Western Blot analysis

Cell lysates were preparing employing RIPA buffer (50mM Tris pH=7.4, 150mM NaCl, 2mM EDTA, 0.5% NP40, 0.25% Sodium Deoxycholate) supplemented with Complete protease inhibitor and PHOSStop phosphatase inhibitor (Roche). Hundred micrograms of protein were separated in 10% SDS PAGE and transferred to a PVDF membrane. The membrane was probed with antibodies for FOXM1, Cyclin B, CHK1 S345, (CST), Rad51, Exo1 (Abcam), RPA2 s4/s8 (Bethyl), GAPDH (GeneTex), tubulin (T6074, Sigma). Bound antibodies were detected with infrared fluorescent secondary antibodies and imaged with a LI-COR Odyssey system. The numbers below the bands are fold of change over the non-treated condition calculated from the relative intensity of the bands quantified employing Image Studio Software.

### Growth in soft agar

HBEC3-KT cells were irradiated in triplicate flasks and transfected at day 4 with the indicated DNA. At day 6, samples were collected to analyze for micronucleus formation. Cells were continuously passaged for 3 weeks. At week 4 media was changed to a 1:1 mixture of complete Keratinocyte Serum Free media and 10% FBS RPMI to accommodate potential metabolic changes and promote clone growth (82). At the end of week 4, 50 000 cells per well were plated in 0.37% agar in triplicate wells over a 0.7% agar bottom layer. Colonies of a diameter larger than 10 cells were counted after 3 weeks.

### Statistical analysis

The statistical test applied in each case is stated in the figure legend. Excel was used for two-tailed Student’s t-test analysis assuming equal variance of the samples. Kaleidagraph (Synergy Software) was used for One Way ANOVA employing a Bonferroni or Fisher’s LSD post-hoc test as indicated in the figure legend. Statistical significance is represented by asterisks: *= p≤0.05; **= p≤0.01; ***= p≤0.005; ****= p≤0.001.

## Supplementary Figures

**Supplementary Figure 1**: Detail of the constructs engineered in U2OS cells to report the repair of a double strand break introduced by cleavage with the endonuclease I-SceI. GFP expression is gained when repair occurs by the specific mechanism.

**Supplementary Figure 2**: Heat Map of the relative expression of all the antigens profiled by the reverse phase protein array after hierarchical clustering. Triplicate samples of non-irradiated (NI) or 2Gy irradiated HBEC3-KT cell lysates collected at day 5 or day 7. Heat map represents ‘‘rank-ordered’’ changes induced by each treatment, calculated by summing median-centered normalized protein amount.

**Supplementary Figure 3**: Characterization of radiation induced replication stress.

A) Asymmetry of replication track: the graph depicts the ratio of CldU/IdU track length for each dose. No statistical divergence following One Way ANOVA analysis.

B) Micronucleus formation rates in HBEC3-KT or U2OS cells were irradiated or treated for 48h with 25μM HU with or without 30μM nucleosides. Error bars are SEM. Student’ t-test. 1 of 2 experiments.

**Supplementary Figure 4**: Correlation of FOXM1 expression with transcriptional targets and replication stress marker levels in cancer cell lines samples analyzed with the RPPA platform:

A) Relative FOXM1 expression across datasets of cell lines grouped by cancer type extracted from the MD Anderson Cell Lines Project Portal. https://tcpaportal.org/mclp/#/

B) Table listing the correlation factor and significance of paired comparison of the indicated protein with FOXM1 expression in each dataset of cell lines grouped by cancer types. Included are the comparisons that were significant (p≤ 0.05).

**Supplementary Figure 5**: Correlation of FOXM1 expression with transcriptional targets and replication stress marker levels in tumor samples analyzed with the RPPA platform:

A) Relative FOXM1 expression across datasets of tumor samples grouped by cancer type extracted from the The Cancer Proteome Atlas. https://tcpaportal.org/tcpa/

B) Table listing the correlation factor and significance of paired comparison of the indicated protein with FOXM1 expression in each dataset of tumor samples grouped by cancer types. Included are the comparisons that were significant (p≤ 0.05)

